# Elucidating the systemic response of wheat plants under waterlogging based on transcriptomic and metabolic approaches

**DOI:** 10.1101/2022.08.03.502608

**Authors:** Geeisy Cid, Davide Francioli, Steffen Kolb, Yudelsy Antonia Tandron Moya, Nicolaus von Wirén, Mohammad-Reza Hajirezaei

**Affiliations:** Leibniz Institute of Plant Genetics and Crop Plant Research (IPK); Leibniz Centre for Agricultural Landscape Research (ZALF)

**Keywords:** Alanine, amino acids, carbohydrates, systemic response, waterlogging, wheat

## Abstract

Extreme weather conditions lead to significant imbalances in crop productivity, which in turn affect food security. Flooding events cause serious problems to many crop species such as wheat. Although metabolic readjustments under flooding are important for plant regeneration, underlying processes remain poorly understood. Here, we investigated the systemic response of wheat to waterlogging using metabolomics and transcriptomics. A 12-day exposure to excess water triggered nutritional imbalances and disruption of metabolite synthesis and translocation, reflected by reduction of plant biomass and growth performance. Metabolic and transcriptomic profiling in roots, xylem, and leaves indicated anaerobic fermentation processes as a local response occurring in roots. Differentially expressed genes and ontological categories revealed that carbohydrate metabolism plays an important role as a systemic response. Analysis of the translocation rate of specific compounds in the xylem showed how waterlogging alters the composition of xylem exudates and thus the root to shoot communication. Interestingly, among all metabolites determined in our study, alanine was the most abundant transported in the xylem. Our results suggest an important role of this amino acid not only as amino-nitrogen source but also as the major root-to-shoot translocated and systemically acting metabolite crucial for balancing C/N between roots and shoots during waterlogging.

The relevance of this study relies on the basis to characterize the important role of alanine as

**Highlight:** Metabolic and transcriptomic changes in wheat highlight alanine as the major root-to-shoot translocated and systemically acting metabolite crucial for balancing C/N between roots and shoots.

## Introduction

Ongoing global warming is one of the current challenges with impacts on weather conditions and serious implications for existing agriculture in terms of productivity, plant morphology and physiology. Among the various climate disturbances, flooding causes detrimental damage to agricultural systems where crop plants such as wheat are exposed to unfavorable growing conditions. It is reported that more than 10% of arable land may be affected by flooding (FAOSTST, www.fao.org). As sessile organisms, plants sense the flooding and adapt their metabolism to survive the prolonged stress, but this is often accompanied by a decline in growth and reproduction. The major consequence of flooding is the partial (hypoxia) or complete (anoxia) depletion of oxygen affecting the central metabolism of plants (Schulze *et al*., 2019; José León 2020). In recent years, tissue-specific investigations have been conducted to understand the morphological, molecular and physiological adaptations of plants in various organs, including roots, shoots or whole seedlings under flooding (Ellis, *et al*., 1999; Kreuzwieser *et al*., 2009; Hsu *et al*., 2011; Mustroph *et al*., 2014; Lothier *et al*., 2020). Among signaling molecules moving from roots to shoots, ethylene has been demonstrated to play a crucial role in the adaptative response of plant organs to hypoxia, including aerenchyma formation, appearance of adventitious roots and epinasty (Schulze *et al*., 2019). It has been reported that inhibitors of 1-aminocylopropane-1-carboxylic acid (ACC), the precursor of ethylene biosynthesis, reduced the synthesis of the latter and thus the epinasty of petioles in tomato plants exposed to waterlogging (Bradford *et al*., 1982). In addition, it has been documented that flooding increases indoleacetic acid (IAA), while abscisic acid (ABA) and jasmonic acid (JA) decreased rapidly (Arbona and Gómez-Cadenas 2008). Among biochemical adaptations, ethanol and lactate production is used by plants to compensate for the losses of glycolytic intermediates and energy in form of ATP to maintain cell viability in the short run. However, alternative pathways and mechanisms are required to maintain a constant flow of assimilates. In this respect, sugars and amino acids, produced in the shoots, were reported to play a major role in translocation and assimilation of the substrates necessary for glycolytic metabolism (De Sousa and Sodek 2003; Lothier *et al*., 2020). An accumulation of soluble sugars, polyols and starch was observed in sunflower leaves (Cui *et al*., 2019) and other plants under waterlogging (Irfan *et al*., 2010). In contrast, inhibition of the delivery rate of specific amino acids e.g., glutamine (Gln) in the xylem of waterlogged soybean plants has been observed (Oliveira *et al*., 2013). Alanine (Ala), Gln, gamma-aminobutyric acid (GABA) and putrescine have often been identified as major amino acids during waterlogging (Reggiani 1999). In this tissue, the transamination of glutamate (Glu) with pyruvate, which is catalyzed by the enzyme alanine aminotransferase (ALAT) induced in rice roots, is thought to be the main source of Ala production (Reggiani 1999; Reggiani *et al*., 2000). Interestingly, high levels of Ala were also found in xylem sap of soybean in hypoxia conditions, suggesting that this amino acid is an important compound translocated from roots to shoots (De Sousa and Sodek 2003). Extensive studies have been focused on the function of Ala production under oxygen deficiency via ALATs (Miyashita and Good 2008; Rocha, Sodek, *et al*., 2010; Bailey-Serres *et al*., 2012). The readily reversible activity of this enzyme and its multiple isoforms has been seen to play an important role on Ala production to conserve nitrogen and carbon that would be lost by ethanol production (Watson *et al*., 1992; Orzechowski *et al*., 1999). However, it remains poorly understood how and to what extent Ala is translocated and further degraded to meet the energy demand in energy-deprived plants. Interestingly, other ALAT homologs with glyoxylate aminotransferase (AGXT) activity in Arabidopsis have been documented in humans using alanine:glyoxylate as substrate to produce pyruvate and glycine (Gly) (Liepman and Olsen 2003). Despite the well documented functions of distinct types of AGXTs in humans, it remains poorly understood what is the function of this enzyme in plants for Ala degradation in scenarios e.g. of flooding which cause higher energy demand. Further clarification is necessary to understand the function of Ala translocation and/or degradation for the recycling of C-N sources to maintain the activity of sugar and/or glycolytic metabolism and to provide a continuous flow for energy production as a systemic response in wheat plants under waterlogging.

In this study, we conducted greenhouse experiments in which wheat plants were exposed to waterlogging for 12 days at tillering stage and subsequently recovered. First, we monitored the physiological changes and nutritional status during and after the stress. Second, we assessed the overall changes in the metabolome and transcriptome that occur in leaves and roots of wheat plants subjected to waterlogging to dissect those related to local or systemic responses. Third, we identified major changes in nutrient, amino acid and hormone translocation rates in the xylem of stressed plants. Based on our findings, we hypothesize that waterlogging triggers systemic responses in wheat. This response is regulated by readjustments in carbohydrate metabolism where translocation of Ala from roots to shoots is essential to provide extra carbon skeletons for *de novo* glucose biosynthesis as extra energy source.

## Materials and Methods

### Experimental set-up

To investigate the metabolic and transcriptomic changes of wheat plants var. Chinese spring exposed to waterlogging stress during tillering stage, a pot experiment referred as **“central experiment”** was conducted in a greenhouse at the Leibniz Institute of Plant Genetics and Crop Plant Research (IPK-Gatersleben, Germany) according to (Francioli *et al*., 2021). Waterlogging condition was conducted in one set of the existing plants for 12 days. After this period, phenotypic traits such as shoot and root fresh and dry biomass, plant height root length and number of tillers were recorded. Fully expanded leaves were quickly collected and stored in liquid nitrogen. Thereafter, roots from the same plants were gently shaken to remove soil particles, carefully washed with tap water and dried with thin towel papers in the less time possible. All samples were immediately stored in -80°C until further analysis. Sampling protocol was conducted in series and completely randomized.

Following the same basic principles in terms of plant developmental stage, duration of the stress and replicate number of the central experiment, a study of the dynamics of growth and metabolism of wheat plants during the 12 days of exposition was performed. The term **“dynamics experiment”** was used to differentiate the results from the previous experiment. In addition to fully developed leaves and roots, xylem exudates were also collected. All plant materials were harvested at different time points, including 0, 3, 6, 9, and 12 days after the onset of waterlogging (Supplementary Fig. S1), followed by a recovery period of 12 days during which excess water was removed from the soil. During this period, the same plant material was harvested at days 2, 6 and 12 from recovered, non-recovered and well-watered plants.

After 12 days of recovery, water excess was removed from the remaining non-recovered plants, and yield parameters e.g. spike and grain weight (g·plant^-1^) and spike and grain number, were monitored in all treatments when plants reached the ripening stage. A final assessment of the germination rate of representative seeds from each treatment was performed.

### Collection of xylem exudates

Xylem exudate was collected at the different time points established in the dynamics experiment previously described. The collection procedure was performed according to (Aguirre 2020) under control conditions at 21°C, 98% humidity and normal light conditions. Eight different biological replicates per time point were used for each treatment. Each biological replicate consisted of three individual plants grown in one pot.

The shoots of the same plants were further used for total fresh and dry biomass and to quickly collect fully expanded leaves for biochemical analyses. Silicon tubes with 2-2.5 mm diameter and 4-5 cm length (Rotilabo®-Silikonschlauch, Carl Roth GmbH + Co. KG, Karlsruhe, Germany) were placed to match the corresponding incision diameter of the hypocotyl and allowed the accumulation of xylem exudates with a total collection time of 4 hours. The silicon tubes were covered with aluminum foil (Rotilabo®-Typ R 100, Carl Roth GmbH + Co. KG, Karlsruhe, Germany) to protect the exudates from contamination and UV light. The estimation of the total volume collected, was registered from the difference in weight of the collection tubes before and after collection.

### Total chlorophyll determination

Total chlorophyll extraction was performed using 10 mg of fresh, fully expanded leaves harvested at day 12 of waterlogging. Samples were incubated in acetone 80% for 60 min and centrifuged at 14 000 rpm for 5 min. Samples were analyzed photometrically at 663 and 645 nm. The quantification of chlorophyll was performed according to Roca *et al*., 2016.

1. Chl a=12.7x(A663) - 2.69x(A645);
2. Chl b=22.9x(A645) - 4.68x(A663);
3. Total Chl=20.2x(A645) + 8.02x(A663)

### The analysis of macro- and micronutrients

Fully expanded leaves and roots (ca. 30 mg) of control and waterlogged treatments from central experiment were weighed into PTFE digestion tubes and concentrated nitric acid (1 mL; 67-69%, Bernd Kraft, Duisburg, Germany) was added to each tube. After 4h incubation, samples were digested under pressure using a high-performance microwave reactor (Ultraclave 4; MLS, Leutkirch, Germany). Digested samples were transferred to Greiner centrifuge tubes and diluted with de-ionized (Milli-Q®) water to a final volume of 15 mL. For the EA in xylem exudates of the same treatments but from the dynamics experiment, 0.005 mL of the samples were directly diluted to a final volume of 5 mL using 5% nitric acid. Elemental analysis was carried out using Inductively Coupled Plasma-Optical Emission Spectroscopy (iCAP 7400 duo OES Spectrometer; Thermo Fisher Scientific, Dreieich, Germany) coupled to an 4DX preFAST automated inline dilution system (Elemental Scientific™ (ESI), Mainz, Germany). A three points external calibration curve was set from a certified multiple standards solution (Bernd Kraft, Duisburg, Germany). Element Yttrium (Y) was used as internal standard for matrix correction.

### Isolation of RNA and preparation for microarray

Total RNA was extracted from fully expanded leaves and roots from central experiment using the NuceloSpin RNA Kit for extraction and NanoDropTM2000c for RNA quality as described in (Beier *et al*., 2022). Library preparation for transcriptome sequencing was performed by Novogene Co. using NEBNext® UltraTM RNA Library Prep Kit for Illumina® (NEB, USA) following manufacturer’s recommendations and considering three biological replicates for each plant material. mRNA was purified from total RNA using magnetic beads bound to poly-T oligos followed by fragmentation. Random hexamer primer and M-MuLV Reverse Transcriptase (RNase H-) were used for first strand cDNA synthesis, whereas second strand cDNA synthesis was performed with DNA Polymerase I and RNase H. cDNA fragments of 150∼200 bp were selected by purification of library fragments using AMPure XP system (Beckman Coulter, Beverly,USA). PCR was performed with Phusion High-Fidelity DNA polymerase, Universal PCR primers and Index (X) Primer. Library quality was assessed using the Agilent Bioanalyzer 2100 system. After cluster generation, the library preparations were sequenced on an Illumina platform and paired-end reads were generated. Reference genome and gene model annotation files were downloaded from genome website browser (NCBI/UCSC/Ensembl) and paired-end clean reads were mapped to the reference genome using HISAT2 software. Feature counts were used to count the number of mapped reads for each gene, including known and novel genes. The RPKM of each gene was then calculated. Differential expression analysis was performed using DESeq2 R package. The resulting P values were adjusted using the approach of Benjamini and Hochberg’s to control the False Discovery Rate (FDR). GO was implemented by clusterProfiler R package and adjusted P value < 0.05 and log2 foldchange ≥ |2| were classified as differentially expressed. For KEGG enrichment analysis, the clusterProfiler R package was used to test the statistical enrichment of differentially expressed genes in KEGG pathways.

### Extraction and quantification of carbohydrates

Soluble sugars and starch were determined according to (Tula *et al*., 2020) with slight modifications. 50 mg of frozen, fully expanded leaves and roots of waterlogged and control plants from central experiment were homogenized in liquid nitrogen, dissolved in 0.7 mL of 80% (v/v) ethanol and incubated at 80°C for 60 min at 600 rpm. The resulting crude extracts were centrifuged at 14 000 rpm at 4°C during 5 min and the supernatant was transferred to a new Eppendorf tube. The remaining pellet was washed two times each 1 ml with 100% ethanol. To break down starch into glucose moieties, 0.2 mL of 0.2 N KOH was added followed by incubation overnight at 4°C and neutralization with 0.07 mL of acetic acid 1N, pH:6.5-7.5. The samples were digested using 0.1 mL of a buffer containing 7 U·mg^-1^ i.e. 2 mg·mL^-1^ amyloglucosidase in 50 mM NaAc, pH:5.2 and incubated overnight at 37°C Starch was calculated from the glucose concentration present in the samples after starch breakdown.Glucose, fructose and sucrose were determined by a coupled photometric assay in the presence of the auxiliary enzyme’s hexokinase, phosphoglucoisomerase, and invertase, respectively, and monitoring the oxidation of NAD to NADH at a maximum wavelength of 340 nm.

### Analysis of primary metabolites

Metabolite extraction followed the protocol described by (Tognetti *et al*., 2007) with some modifications. 100 mg of frozen leaves and roots from central experiment were used. Samples were homogenized thoroughly at 4°C with gentle shaking for at least 20 min. Subsequently, 0.3 mL HPLC grade water was added to each sample and centrifuged at 14.000 rpm at 4°C for 10 min. The supernatant was transferred to a new Eppendorf tube and dried in a Speed-Vac (Christ RVC2-33IR, Germany) at 35°C for approximately 2 hours. The resulting pellet was resuspended in 0.3 mL HPLC grade water and immediately used for the measurement of desired compounds. The separation and detection of metabolites was performed according to (Ghaffari *et al*., 2016), using an ion chromatography system connected to a conductivity detector (Dionex, Thermofisher Germany) and a triple quadrupole Mass Spectrometer QQQ6490 (Agilent Technologies, Waldbronn Germany). The gradient was produced from ultrapure water (buffer A, Millipore) and a concentrated potassium solution EGCIII KOH (Dionex, Germany, buffer B) using an eluent generator EG-SP (Dionex Germany). The column was equilibrated at the same conditions at 0.32 mL·min-1 and 35°C throughout the measurement. ESI-MS/MS analysis was conducted in negative ionization mode using nitrogen gas at 720 L·h^-1^ at a heating temperature of 250°C, capillary voltage 3.5 KV and different dwell times between 40 and 200 sec. Collision energy (CE) ranged from 6 and 50 for different masses. Multiple reactions monitoring (MRM) was performed to accurately identify individual compounds. Quantification was performed considering 30 different compounds and a calibration curve of the same with concentrations in the range between 25 and 500 pmol·µL-1. Data acquisition and quantification were performed using Chromeleon software, release 7.3 (Dionex GmbH, Germany) and MassHunter software, release B.07.01 (Agilent Technology, Waldbronn Germany).

### Determination of hormone profile in xylem exudates

For phytohormone analysis in xylem exudates, 0.03 mL of the samples were mixed with internal standards (OlChemim s.r.o, Czech Republic) at a final concentration of 100 nM. Samples were homogenized 30 sec and centrifuged 30 min at 14000 rpm and 4°C (CT 15 RE centrifuge, Himac, Japan). The analysis of auxins and cytokinins was performed by UHPLC-ESI-MS/MS as described in (Eggert and von Wirén 2017).

Gibberellins were determined using UHPLC-HESI-HRMS in negative ion mode. A reversed phase Acquity UPLC HSS T3 column (10 Å, 2.1×150 mm, 1.8 μm, Waters) (45°C, 0.3 mL·min^-1^) coupled to a guard column (130Å, 2.1×5 mm, 1.8 μm, Waters) was used. The gradient consisted of A (Water, 0.1% FA) and B (MeOH, 0.1% FA) as follows: 0–0,3 min, 10% B; 0,3–0,7 min, 10% to 30% B; 0,7–2 min, 30% to 50% B; 2–4 min, 50% to 60% B; 4–8 min, 60% to 80% B; 8–9,5 min, 80% to 99% B; 9,5–10,4 min, 99% B. Source values were set as follow: Spray voltage 2.5 kV; capillary temperature 255°C; S-lens RF level 40; Aux gas heater temp 320°C; sheath gas flow rate 47; aux gas flow rate 11. The spectra acquisition was performed on a Full MS/dd-MS^2^ experiment. Resolution in Full Scan was set as 70000. Resolution of the MS/MS experiments were 17,500 and NCE 40V was used. Data acquisition and processing were performed using Trace Finder Software (v. 4.1, Thermo Scientific, San Jose, CA, USA). To generate the calibration curve, the peak area was measured on the extracted ion chromatogram (XIC) of the de-protonated molecule ion [M-H]^-^. Compounds were identified based on their retention times, high resolution m/z spectrum and isotope pattern in respect to standards. Additionally, generated MS2 spectra were searched in a custom spectral library for the confirmation of compound identification.

### Amino acids extraction and quantification

For the analysis of amino acids in leaves and roots from central experiment, the same extracts were used as for sugar analysis. The translocation rate of the same amino acids in xylem exudates of plants from dynamics experiment was also analyzed. Plant tissue or xylem sap were derivatized using an NMR-purified fluorescing reagent AQC (6-aminoQuinolyl-N hydroxysuccinimidyl carbamate) (BIOSYNTH AG, Switzerland). Three mg of AQC were dissolved in 1 ml acetonitrile and incubated for 10 min at 55°C. Derivatization was performed for 0.01 ml of plant extract/ xylem exudate/ standard mixture using 0.01 mL of AQC and 0.08 ml of boric acid 0.2M, pH:8.8 at 55°C for 10 min. Separation of soluble amino acids was performed by ultra-pressure reversed-phase chromatography (UPLC) Acquity H-Class (Waters GmbH, Germany) coupled to fluorescent detector (FLR Detector). Separation was carried out on a C18 reversed-phase column (Luna Omega, 1.6 µm, 2.1×100 mm, Phenomenex, Germany) with a flow rate of 0.6 ml·min^-1^ during 6 min. The column was heated at 40°C during the whole run. The gradient was accomplished using an eluent A concentrate, eluent B and eluent C with purest water according to the manufacture’s manual (Bioanalytics Gatersleben, http://www.bioanalytics-gatersleben.de). The detection wavelengths were 266 nm for excitation and 473 nm as emission. Quantification of amino acids in the samples followed a calibration curve of a mixture of 20 different amino acids in the range 0.5 to 20 nmol·mL^-1^ considering R2> 0.98 in all cases.

### Statistical Analysis

Statistical analyses were performed using R Studio version 4.1.2. Normal distribution of the data was checked using Shapiro-Wilk’s normality test. To determine the difference between paired groups, Student T-test or one-way ANOVA was performed. Different letters indicate significant differences between the treatments based on Tukey’s HSD. In all cases, 95% confidence interval (p≤ 0.05) was considered (*p<0.05; **p<0.01; ***p<0.001; ns: not significant). Heat Map was generated using the online versatile matrix visualization and analysis software MORPHEUS, https://software.broadinstitute.org/morpheus.

## Results

### Influence of waterlogging on different phenotypic traits in wheat plants at tillering stage and yield parameters at ripening stage

Considering the data collected from the dynamics experiment, in which plants were recovered from waterlogging by removal of excess water (see Material and Methods, experimental setup for more details), no significant changes in shoot biomass were observed until day 6 after setting up waterlogging (Fig. 1A, green color). However, a gradual decrease was observed compared to control plants until day 12. After stress removal, plants progressively recovered, reaching the level of control plants at day 12. However, clear differences between control and non-recovered plants were observed form day 6 after removing the stress (Fig. 1B top). Recovered plants exhibited similar height and leaf abundance as the control plants, suggesting the reversible effect of waterlogging on shoot biomass even when the stress was prolonged for several days.

**Fig. 1.**
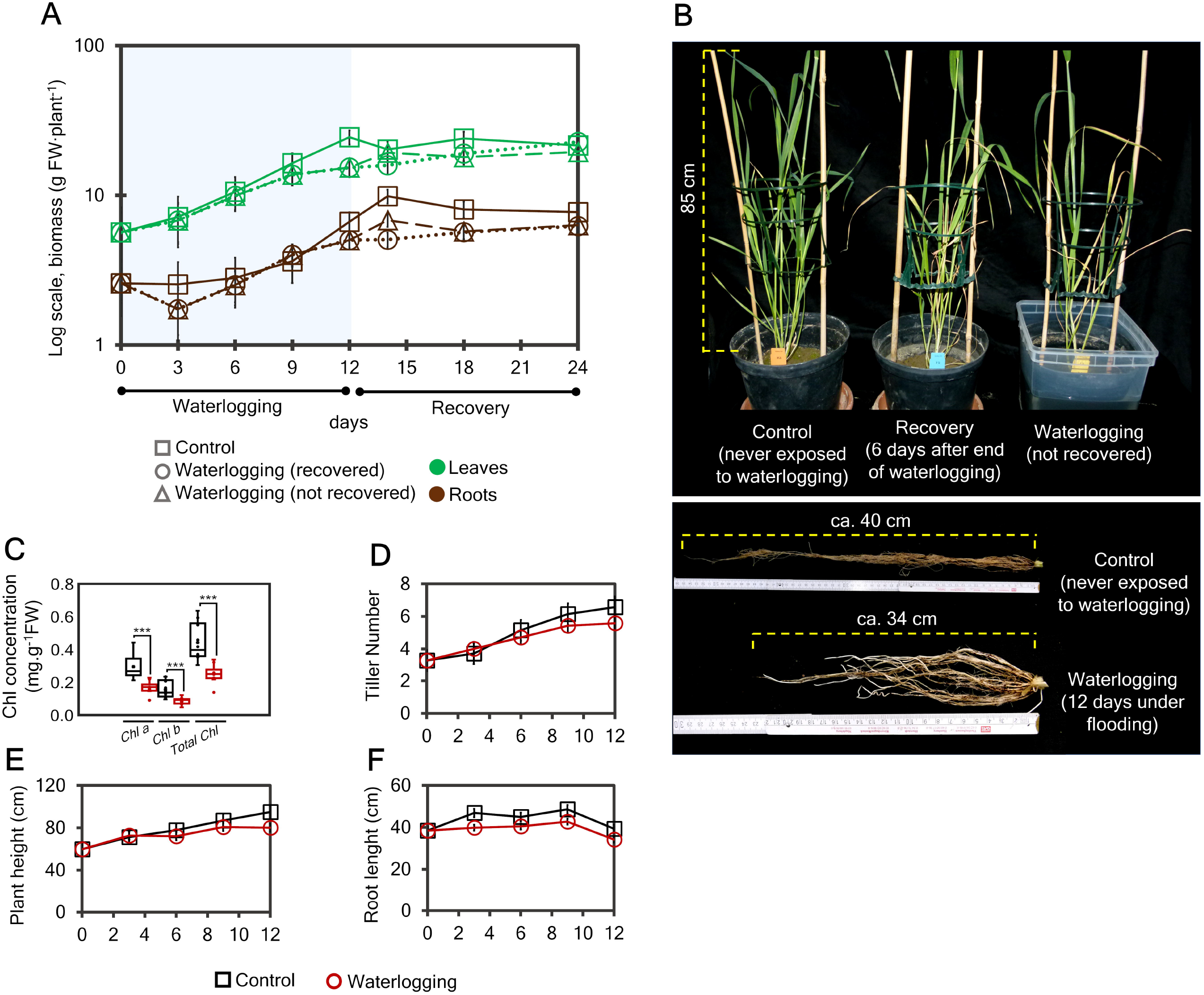
Assessment of phenotypic changes on wheat plants during waterlogging. (A) Dynamics of shoot and root biomass in wheat plants during waterlogging and the recovery period. (B) appearance of shoots and roots of wheat plants under the different treatments. (C) Chlorophyll concentration in fully expanded leaves, (D) Number of tillers, (E) plant height and (F) root length. After the setting up of waterlogging, fully expanded leaves and roots were collected from the control and waterlogging treatments. Different time points were considered during the stress (0, 3, 6, 9 and 12 days under waterlogging) and after removing the excess of water (2, 6 and 12 days after end of waterlogging). Each time point indicates the mean ± SE (n=8) and the bars indicate the mean ± SE (n=10). Normal distribution of the data was checked using Shapiro-Wilk’s normality test. To determine the difference between paired groups, Student T-test (p≤ 0.05) was performed. Asterisks represent ***p<0.001.

At day 12, the photosynthetic capacity of waterlogged plants decreased as total chlorophyll in the leaves was reduced by about twofold compared to the control (Fig.1C). However, the ratio Chl a/b remained unaffected, suggesting a readjustment of the plants to optimize light absorption and photosynthesis under the stress. Other aboveground parameters, such as the dynamics of tiller number and plant height (Fig.1D, E), remained unaffected until day 6 of the stress with non-significant decrease from this time point until day 12. In contrast, the root biomass responded earlier during the exposure time (Fig. 1A, brown color). In waterlogged roots, biomass decreased by 32% on day 3 of exposure. Interestingly, from day 6 up to day 9, roots biomass of stressed plants did not differ to that from control and the appearance of adventitious roots was observed (Fig. 1B bottom). Subsequently, this parameter dropped again from day 9 throughout the recovery period. Root length was also reduced in plants exposed to waterlogging, averaging about 34 cm compared to 50 cm in control plants (Fig. 1F). At ripening stage, the total weight and number of spikes and total weight of grains did not differ in plants recovered from 12 days of waterlogging to that from control (Supplementary Fig. S2). However, the same parameters were decreased in average twofold in plants that did not recover. The total amount of grains counted on recovered and non-recovered plants differed significantly to those counted in control. Recovered plants produced 20% less grains than the control plants, whereas non-recovered plants produced two times fewer grains (Supplementary Fig. S2).

### Effect of waterlogging on hormone translocation in wheat plants

The dynamics of hormone translocation rate in xylem exudates was evaluated from plants of all treatments. Of all hormones determined in this study, gibberellins accounted for the greatest amount and diversity in the xylem (Fig. 2A). The translocation rate of GA53, GA44, GA19, GA5, GA15, and GA4 increased two- or threefold and reached a maximum on the third day of the dynamic experiment in both control and waterlogged plants, followed by a decrease in the following days of stress. Only GA51 decreased steadily from day 0 until day 9. During the recovery phase, the levels of all GA’s remained low in control and non-recovered plants, whereas an increase in average about twofold was observed at day 6 after recovery in those plants where the stress was removed.

**Fig. 2.**
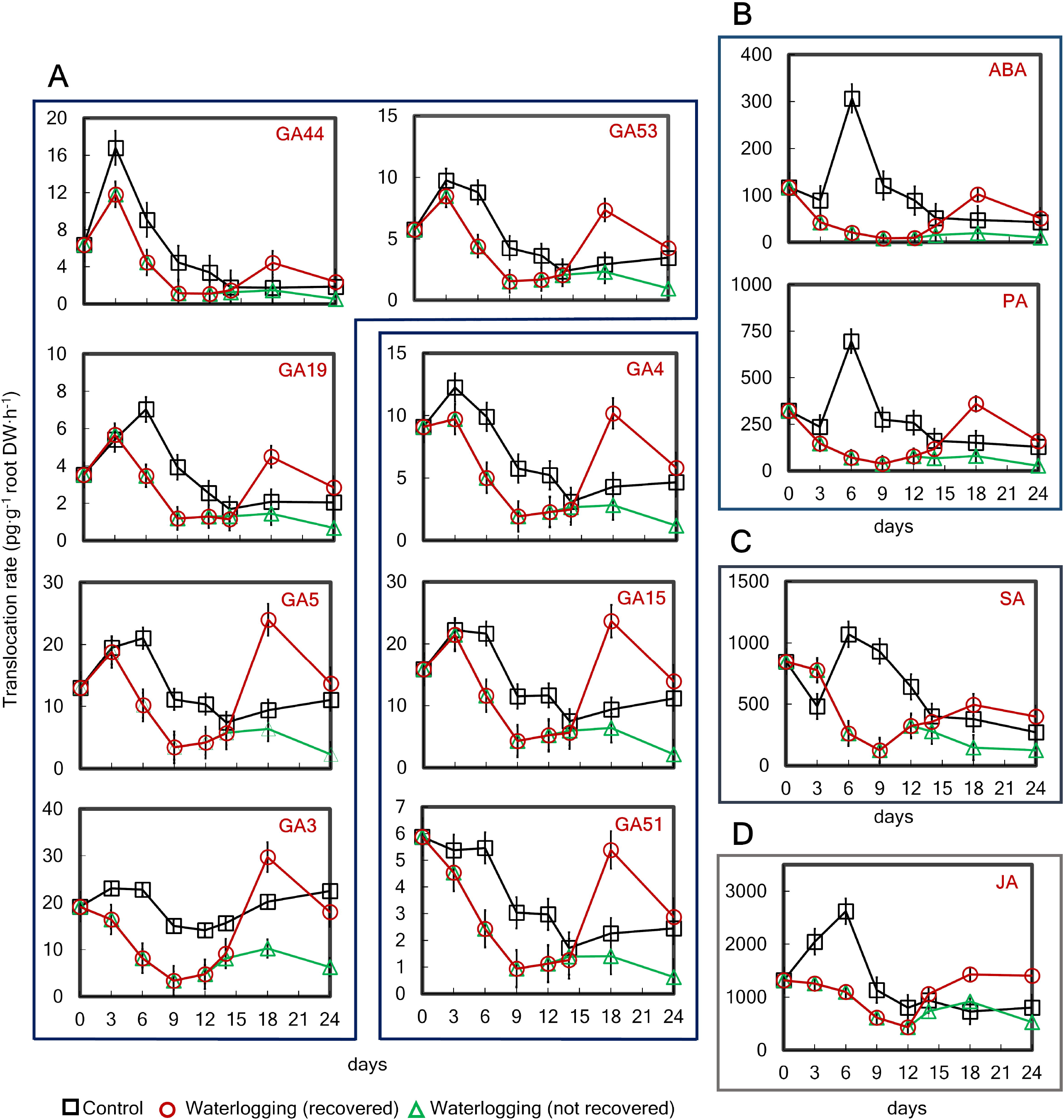
Influence of waterlogging on the hormone translocation rate in the xylem of wheat plants. Dynamics of translocation rate of (A) gibberellins (GA44, GA19, GA5, GA3, GA53, GA4, GA15, GA51, (B) abscisic acid/ phaseic acid (ABA/PA), (C) salicylic acid (SA) and (D) jasmonic acid (JA). After the setting up of waterlogging, xylem exudates were collected from the control and waterlogging treatments. Different time points were considered for collection during the stress (0, 3, 6, 9 and 12 days under waterlogging) and after removing the excess of water (2, 6 and 12 days after end of waterlogging). Each time point indicates the mean ± SE (n=8).

Abscisic acid (ABA) and its degradation product, phaseic acid (PA), were also found in xylem sap (Fig. 2B). Their translocation rate in control plants increased by threefold on day 6 of exposure followed by a decline on day 9. Stressed plants showed a steady decrease until day 12, as observed for gibberellins during the recovery phase. Interestingly, translocation of salicylic acid (SA) increased twofold at day 3 in waterlogged plants followed by a sharp decrease until day 9 (Fig. 2C). Non-recovered plants maintained low SA concentrations, with no significant differences between the time points. In contrast, recovered plants increased SA level to control levels once the stress was removed. A similar pattern to ABA and PA was found for jasmonic acid (JA). Stressed plants decreased the translocation rate of JA from day 0 compared to the control, followed by a twofold recovery compared to the control plants (Fig. 2D).

### Influence of waterlogging on the composition of macro- and micro-nutrients in leaves and roots of wheat plants

The analysis of macro- and micro-nutrient composition was performed in fully expanded leaves and roots exposed 12 days to waterlogging and in xylem exudates collected during the exposure period. Plants under waterlogging showed a reduction of nitrogen (N) concentration in leaves by about 35%, while no significant difference was observed in roots compared to control (Fig. 3A). In contrast, the total carbon (C) concentration increased significantly in leaves and roots compared to control (Fig. 3B). Potassium (K), phosphorus (P), sulfur (S) and zinc (Zn) were reduced in leaves by about twofold, whereas manganese (Mn) increased in the same ratio (Fig. 3C). Other nutrients such as calcium (Ca) and iron (Fe) remained unaffected. In contrast, the concentration of K and P did not vary in roots respect to control, whereas magnesium (Mg), Ca, S and Zn were significantly decreased. Mn was also accumulated in roots from stress plants by about fourfold (Fig. 3D).

**Fig. 3.**
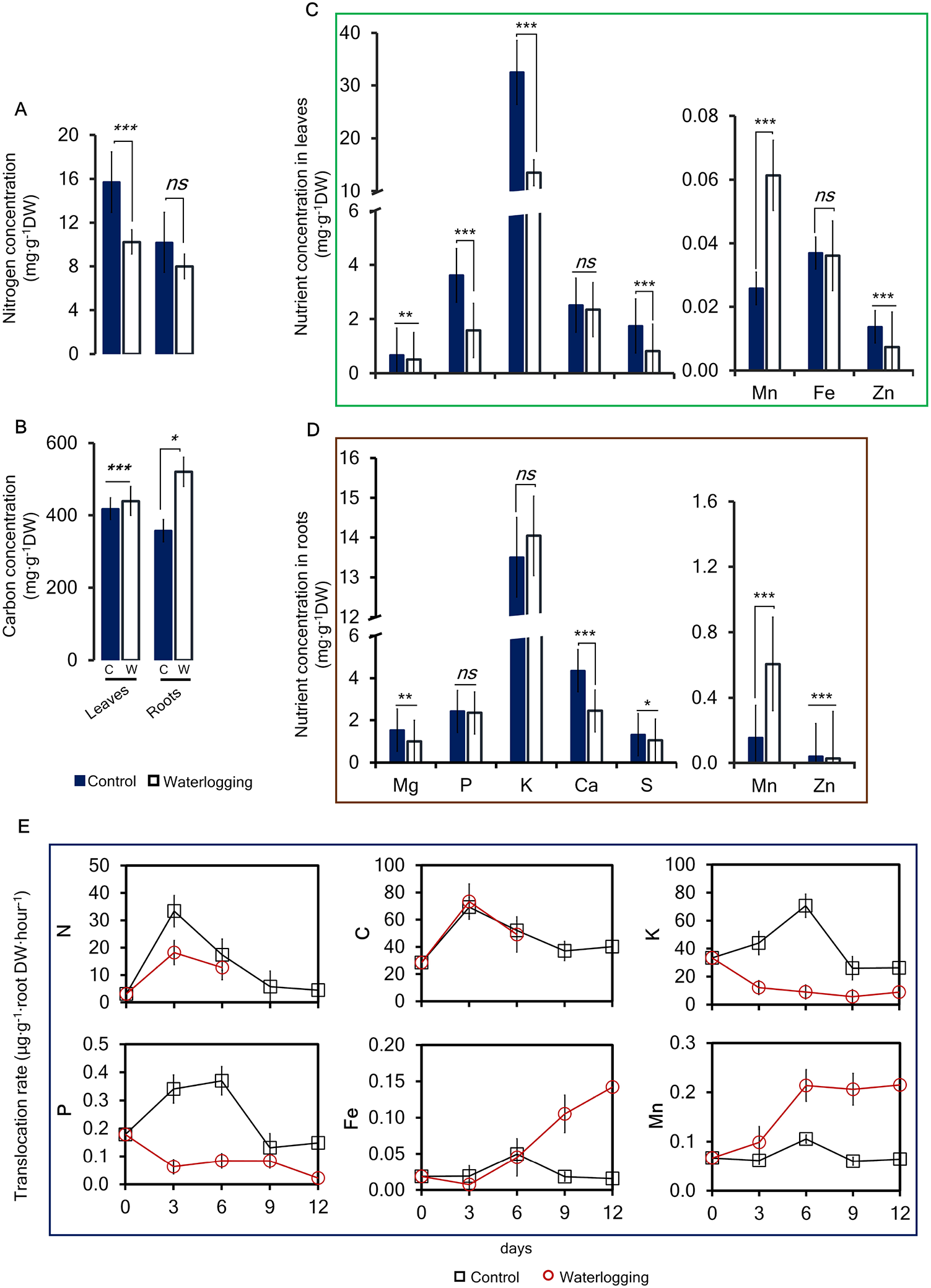
Influence of waterlogging on the mineral composition in fully expanded leaves and roots of wheat plants. (A) nitrogen concentration, (B) carbon concentration and other macro- and micro-elements in (C) fully expanded leaves and (D) roots of control (C) and waterlogging (W) treatments. (D) translocation rate of N, C, K, P, Fe and Mn in the xylem of wheat plants. After the setting up of waterlogging, fully expanded leaves, roots and xylem exudates were collected from the control and waterlogging treatments. Nutrient composition in leaves and roots, are only represented at time point 12 days under waterlogging. Nutrient translocation rate in the xylem was evaluated at different time points during the stress (0, 3, 6, 9 and 12 days under waterlogging). Bars and time points indicate the mean ± SE (n=8). Normal distribution of the data was checked using Shapiro-Wilk’s normality test. To determine the difference between paired groups, Student T-test (p≤ 0.05) was performed. Asterisks represent as follow: *p<0.05; **p<0.01; ***p<0.001; ns: not significant.

The dynamics of the translocation rate of the same elements were measured in the xylem exudates (Fig. 3E). Indeed, N translocation decreased from early time point in stressed plants, whereas C concentration did not differ between control and stressed plants. K and P decreased sharply from day 0, with the highest difference occurring at day 6, where they decreased eight- and fivefold, respectively. In contrast, Mn and Fe were increased in xylem exudates from day 3 and 6, respectively. By day 12 under waterlogging, they increased 4- and 7-fold, respectively, in stressed plants.

### Effect of waterlogging on differentially expressed genes in leaves and roots of wheat plants

To compare the transcriptomic responses between above- and below-ground organs, we first distinguished those metabolic processes that occurred in leaves, roots and in common. Representation of genes filtered at fold change 2 and p-value < 0.05 in a Venn diagram (Fig. 4A) showed that indeed, the major number of genes annotated in leaves were up-regulated (1482), whereas they were down-regulated in roots (4276). Common genes represented in a heat map (Fig. 4B) showed that the up-regulated genes were mainly expressed in leaves, whereas in roots most genes were down-regulated.

**Fig. 4.**
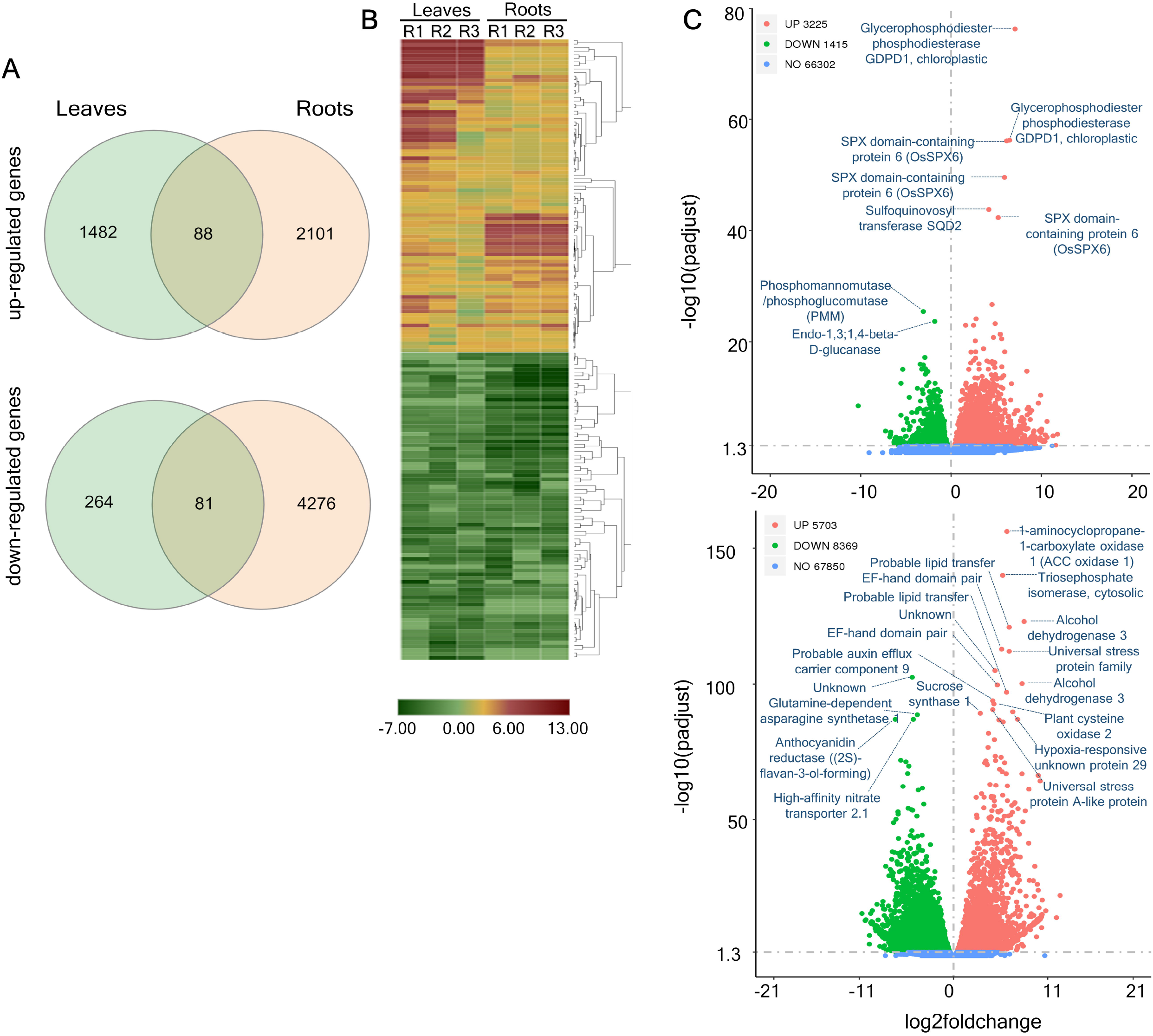
Effect of waterlogging on differentially expressed genes in fully expanded leaves and roots of wheat plants. (A) Venn diagram showing DEGs in fully expanded leaves, roots and in common organs. (B) Heatmap of relative gene expression changes in fully expanded leaves compared to roots of the same plants exposed to waterlogging. C: Volcano plots for the distribution of DEGs in fully expanded leaves (above) and roots (below) from waterlogged plants versus control. In all cases, genes were grouped according to their transcript abundance at day 12 after setting up the waterlogging treatment. Genes were selected considering a threshold >2 in absolute value and padjust < 0.05.

Global gene expression analysis in leaves or roots only compared to the corresponding controls discovered that *glycerophosphodiester phosphodiesterase* (*GDPD1*) (p=0.001), SPX domain-containing protein 6 (p=0.001) and *inorganic pyrophosphatase 2* (*PPi phosphatase 2*) (p=0.001) were highly overrepresented in leaves only (Fig. 4C), whereas genes in the category of core hypoxia response, including *1-aminocyclopropane-1-carboxylate oxidase 1* (*ACC Oxidase*) (p<0.001), hypoxia-responsive unknown protein 29 (p<0.001) and *alcohol dehydrogenase 3* (*ADH3*) (p<0.001) were up-regulated in roots only and not in leaves. Subsequently, common gene expression in both tissues showed up-regulated genes associated to *purple acid phosphatase 23* (*PAP23*) (leaves: p<0.001; roots: p<0.01), SPX domain-containing protein 3 (*OsSPX3*) (leaves: p<0.001; roots: p<0.01), *AP2/ERF* and B3 domain-containing protein (leaves: p<0.001; roots: p<0.001) and *sulfoquinovosyl transferase SQD2* (leaves: p<0.001; roots: p<0.05).

In contrast, expression of genes related to *phosphomannomutase/phosphoglucomutase* (*PMM*) (p<0.01) and *endo-1,3; 1,4-beta-D-glucanase* (p<0.01) were represented in leaves only, as most down-regulated, whereas genes encoding a *probable auxin efflux carrier component 9* (p<0.001), *glutamine-dependent asparagine synthetase 1* (p<0.01), *anthocyanidin reductase (2S)-flavan-3-ol-forming* (p<0.01) and *high affinity nitrate transporter 2*.*1* (p<0.01) were down-regulated in roots only (Fig. 4C). In general, transcriptome analysis revealed that gene expression in both tissues were distinctly regulated under waterlogging and hypoxia responsive genes responded in roots only.

Differential expression of genes related to nutrient transport and assimilation was very diverse among genes annotated in both plant material. For example, genes related to nitrate transporters such as NRT2.1 (p<0.01) and NRT2.4 (p<0.01) were strongly down-regulated in waterlogged roots (Supplementary Fig. S3). Only one gene encoding a NRT2.1 (p<0.01) out of the 20 registered genes was about fivefold up-regulated in stressed roots. Nitrate reductase (NAD(P)H) (p<0.01), high-affinity nitrate transporter accessory (NAR2.1) (p<0.01), ammonium transporter 3.1 (AMT3.1) (p<0.01) and ferredoxin-nitrite reductase 1 (NIR1) (p<0.01) in addition to 7 out of 13 significant total phosphate transporters (PT1/2/4) (p<0.01) and 16 out of 20 potassium-related genes were down-regulated in roots.

Highly up-regulated genes in roots were found to be related to NRT1/PTR family (p<0.01), i.e. CHL1/NRT1.1 (NPF6.3) (p<0.01) and nitrate excretion transporter 1, NAXT1 (NPF2.7) (p<0.01). In addition, genes encoding an inorganic phosphate transporter 1-2 (p<0.01), SODIUM POTASSIUM ROOT DEFECTIVE 2 genes (p<0.1) encoding a heavy metal-binding protein, two Pore potassium Channel b (p<0.01) associated with calcium-activated outward-rectifying potassium channel 2 and Fe2+ transport protein 2 (IRT2) (p<0.001), were also strongly up-regulated in roots. In contrast, all genes related to nutrient transport and assimilation annotated in leaves of stressed plants were highly up-regulated (Supplementary Fig. S3).

### Classification of metabolic pathways involved in leaf and root responses under waterlogging

To uncover metabolic pathways associated to leaf and/or root responses to waterlogging compared to their corresponding controls, KEGG enrichment analysis was performed considering p<0.05 (Fig. 5A). Genes involved in the functional categories of linoleic acid (p<0.01) and amino sugars/nucleotide sugar metabolisms (p<0.05) were significantly higher in leaves only, with the highest number counted as up-regulated genes. Three out of the five significantly regulated genes annotated in the first category, encoded putative *8-, 5-* and *2*.*1-lipoxygenases* (p<0.01), whereas the second category was overrepresented mostly by three out of 12 genes related to pathogenesis-related (*PR)3 chitinases* (two chitinases 4, p<0.01 and p<0.05 and chitinase 8, p<0.01) and other two *UDP arabinopyranose mutase 1*, p<0.01 and *UDP-glucose-6-dehydrogenase 4*, p<0.01) involved in the biosynthesis of cell wall polysaccharides. A wide range of metabolic processes such as glycolysis/gluconeogenesis (p<0.001), amino acids (p<0.001), starch and sucrose metabolism (p<0.01) and plant hormone signal transduction (p<0.001) were represented in roots only being mostly down-regulated. However, known hypoxia-related genes such as *alcohol dehydrogenase 3* (p<0.01), *L-lactate dehydrogenase A* (p<0.01), *ACC oxidase* 1 (p<0.01) and *ACC synthase* 1 (p<0.01) were strongly up-regulated.

**Fig. 5.**
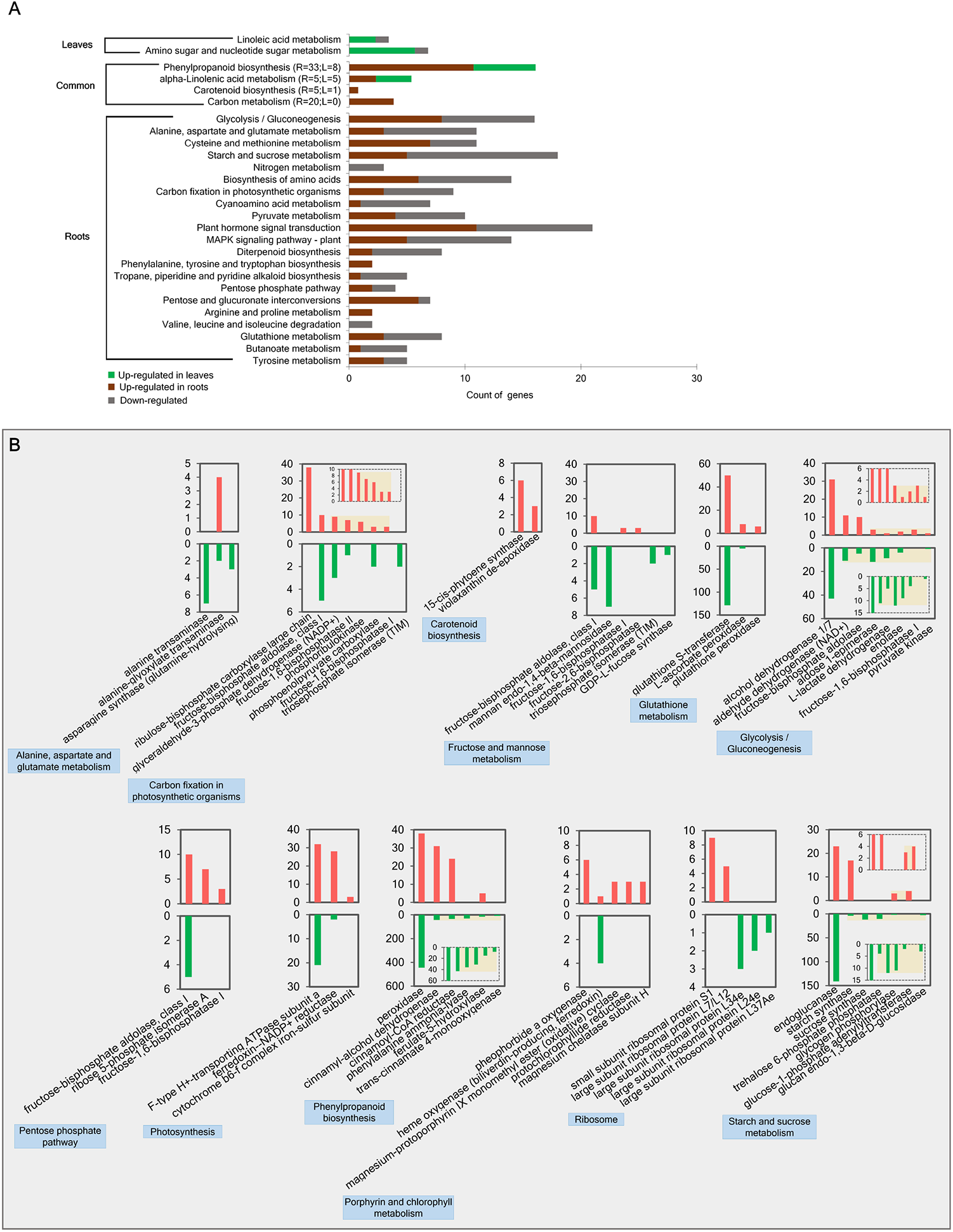
Kyoto Encyclopedia of Genes and Genomes (KEGG) enrichment analysis in fully expanded leaves and roots of wheat plants under waterlogging. (A) Number of genes for metabolic pathways found in fully expanded leaves, roots and in common organs compared to their respective controls. (B) Number of genes for common metabolic pathways found in fully expanded leaves compared with roots of the same plants. In all cases, genes were counted for each metabolic pathway according to their transcript abundance at day 12 after setting up the waterlogging treatment. Genes were counted in each metabolic pathway if their fold-change of expression was >2 in absolute value and padjust < 0.05.

Common metabolic pathways resulting from comparison between leaves or roots with their respective controls were phenylpropanoid (leaves: p<0.05; roots: p<0.01), alpha-linolenic acid (leaves: p<0.01; roots: p<0.05), carotenoid (leaves: p<0.05; roots: p<0.01) and carbon metabolism (leaves: p<0.05; roots: p<0.01) in order of significance (Fig. 5A). However, a better dissection of common metabolic process by comparing leaves and roots from the same stressed plants was also annotated (Fig. 5B). As most significant metabolic processes, considering global p values < 0.05, phenylpropanoid biosynthesis (p<0.001), photosynthesis (p<0.001), starch and sucrose metabolism (p<0.001), glycolysis-gluconeogenesis (p<0.001) and carbon fixation in photosynthetic organisms (p<0.001) were the most significant categories between leaves and roots of the same stressed plants. Consecutively, alanine, aspartate and glutamate metabolism (p<0.001), glutathione metabolism (p<0.001) and fructose and mannose metabolism (p<0.01) were predominant.

Considering the top 50 up-regulated genes in leaves compared to the roots, *glyceraldehyde-3-phosphate dehydrogenases* (*NADP+*) (EC. 1.2.1.13) (p<0.001), *ribulose-bisphosphate carboxylase large chain* (EC. 4.1.1.39) (p<0.001) and *fructose-bisphosphate aldolase, class I* (EC. 4.1.2.13) (p<0.001) were the most overrepresented genes in leaves. In addition, *alanine-glyoxylate transaminase* (EC. 2.6.1.44) (p<0.001), *cytochrome b6-f complex iron-sulfur subunit* (EC. 7.1.1.6) (p<0.001), *fructose-1,6-bisphosphatase II* (EC. 3.1.3.37) (p<0.001), *glutathione S-transferase* (EC. 2.5.1.18) (p<0.001) and *fructose-2,6-bisphosphatase* (EC. 3.1.3.46) (p<0.001) were also higher up-regulated in leaves than in roots. The top 50 most down-regulated genes in leaves corresponded to *alcohol dehydrogenase 1/7* (EC. 1.1.1.1) (p<0.001), *peroxidase* (EC. 1.11.1.7) (p<0.001), *endoglucanase* (EC. 3.2.1.4) (p<0.001), *sucrose synthase* (2.4.1.3) (p<0.001), *phenylalanine ammonia-lyase* (EC. 4.3.1.24) (p<0.001).

### Effect of waterlogging on the metabolic profile of leaves and roots in wheat plants

Key metabolites from glycolysis and TCA cycle were analyzed in fully expanded leaves and roots of plants exposed to 12 days to waterlogging (Fig. 6). Interestingly, sugars accumulated in both, leaves and roots of waterlogged plants (p<0.01), with exception of sucrose (Fig. 6A) and hexose-phosphates (glucose-1-phsopahte: fructose-6-phosphate: glucose-6-phosphate) (Fig. 6E) that did not change in leaves. The concentration of 3-phosphoglyceric acid (3PGA) (F) (p<0.01) was found to significantly decrease in leaves of waterlogged plants, whereas it was not detected in roots of the same plants. Pyruvate (Fig. 6G) was strongly reduced in leaves (p<0.001), whereas it did not change in roots. In contrast, lactate (Fig. 6H) was not detected in leaves and did not change in roots of stressed plants. Other intermediates of respiratory metabolism such as citrates (iso-citrate, citrate) (Fig. 6I) (p<0.01), succinate (p>0.05) (J) and 2-oxoglutarate (Fig. 6K) (p<0.01) were decreased or not changed in leaves and accumulated in roots (p<0.01).

**Fig. 6.**
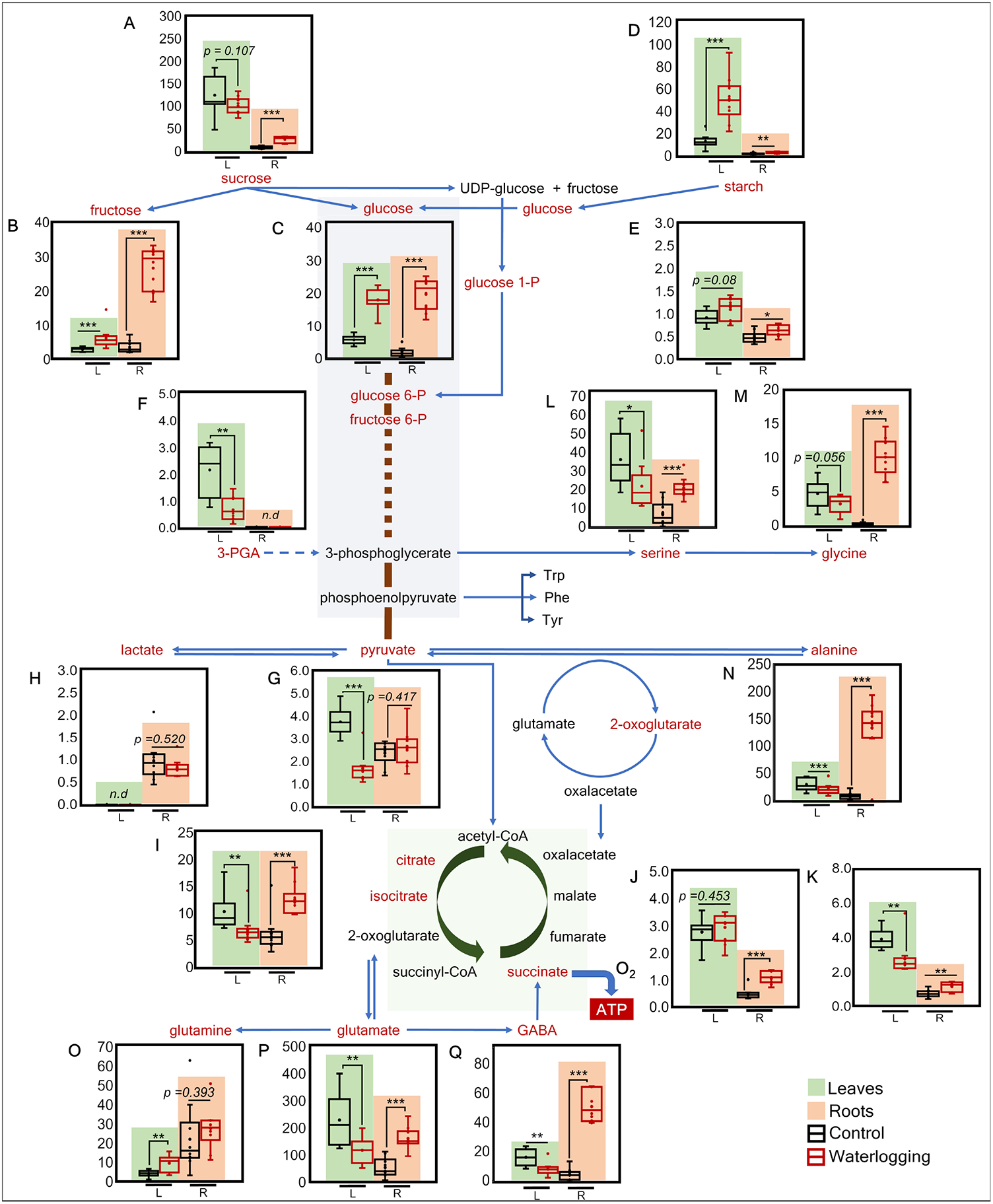
Changes of the metabolite profile in leaves and roots of wheat plants under waterlogging. Concentration of soluble sugars, starch, organic acids and amino acids are represented in fully expanded leaves (green background) and roots (orange background). The metabolites measured in both organs are given as follows: (A) sucrose, (B) fructose, (C) glucose, (D) starch, (E) hexose-P, (F) 3-PGA, (G) pyruvate, (H) lactate, (I) citrates (iso-citrate/citrate), (J) succinate, (K) 2-oxoglutarate, (L) serine, (M) glycine, (N) alanine, (O) glutamine, (P) glutamate, (Q) gamma-aminobutyric acid. After 12 days of setting up waterlogging, fully expanded leaves and roots were collected from the control and waterlogging treatments. Sucrose, glucose, fructose and starch are expressed in µmol·g^-1^ FW, the remaining compounds in nmol·g^-1^ FW. The line in the middle of the boxes are plotted as the median and box limits indicate the 25^th^ and 75^th^ percentiles. Whiskers extend to 1.5 times the interquartile range from the 25^th^ and 75^th^ percentiles. Normal distribution of the data was checked using Shapiro-Wilk’s normality test. To determine the difference between paired groups, Student T-test (p≤ 0.05, n=8) was performed. Asterisks represent as follow: *p<0.05; **p<0.01; ***p<0.001.

Changes in amino acids concentration were observed in stressed plants (Supplementary Table S1). In general, a decrease in the amino acid concentration was observed in leaves, whereas an accumulation of the same was found in roots. Only methionine (Me) (p<0.001) and glutamine (Gln) (p<0.01) accumulated in leaves by about fourfold and twofold, respectively. However, the most significant changes under waterlogging occurred for glycine (Gly) (p<0.001), alanine (Ala) (p<0.001 and gamma-aminobutyric acid (GABA) (p<0.001), which increased by about 26-, 20- and 11-fold, respectively. Serine (Ser) (p<0.001) and glutamate (Glu) (p<0.001) were also increased in roots by threefold. Interestingly, only asparagine (Asn) (p<0.05) decreased fivefold in stressed roots. The same amino acids were determined in xylem exudates during the 12 days under the stress and recovery phase. In contrast to what was found in leaves and roots, the translocation rate of most amino acids was lower in stressed plants. However, a high translocation rate was registered for Ala (p<0.001), GABA (p<0.001) and Gly (p<0.001) along all exposure times reaching the highest translocation between day 3 and day 6. At day 6 after removing the stress, the same amino acids reached similar values to those of control plants. Interestingly, of all metabolites determined in xylem exudates, Ala accounted for the most abundant under waterlogging during the 12 days of exposition (Fig. 6N).

Performing a principal component analysis (PCA) considering the normalized data of 28 metabolites and eight different elements quantified in this study, allowed to discriminate their profile between roots and leaves in PCA1 (Fig. 7). The differences between waterlogging and control in both tissues were explained in PCA2. Considering the length and direction of the vectors, the distributions of the loading plots confirmed that fructose, Ala, Mn and to a lesser extent glucose, strongly influenced the cluster formed by waterlogged roots. Interestingly, among these variables, alanine and glucose vectors showed the highest positive correlation with each other.

**Fig. 7.**
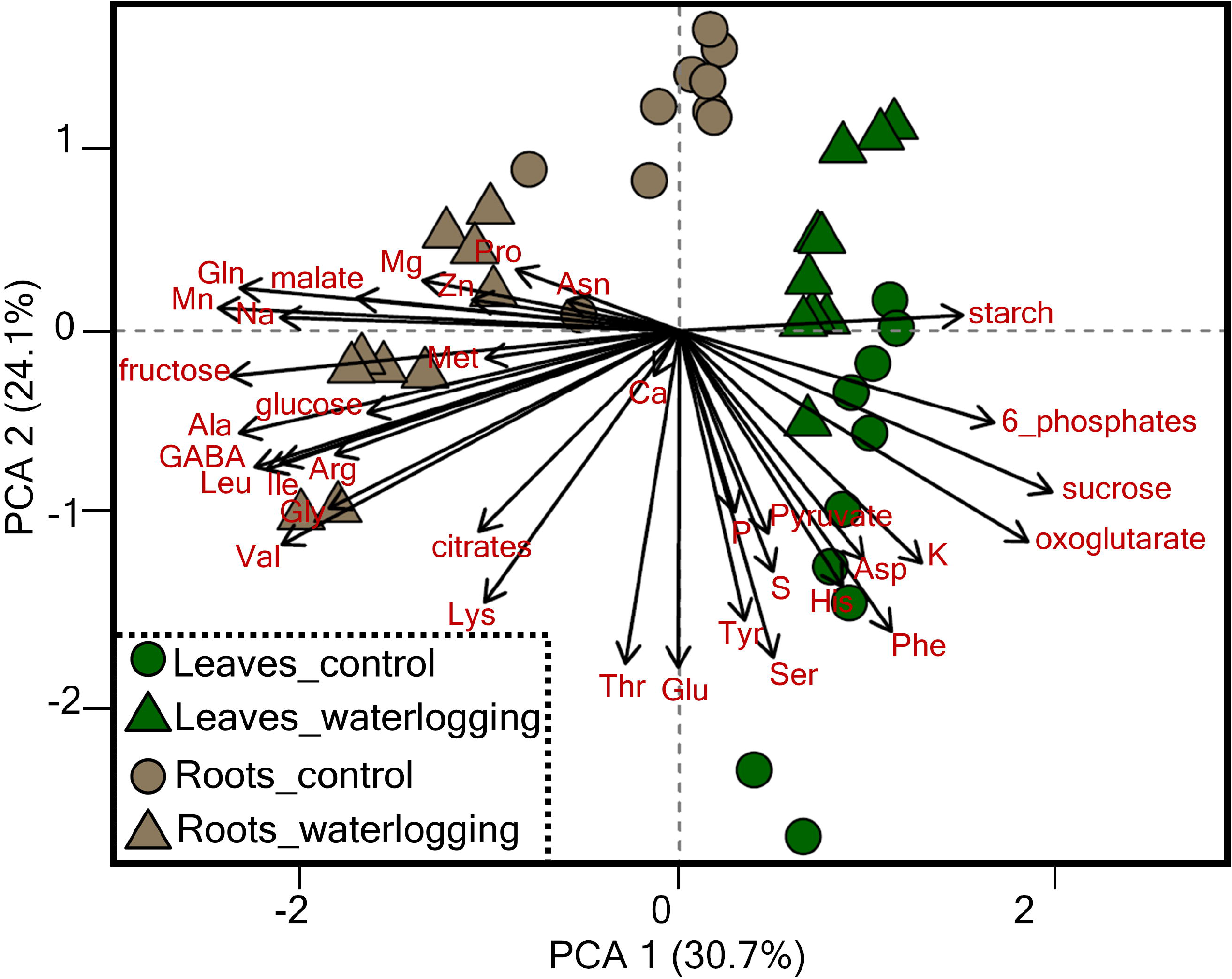
Principal Component Analysis-Biplot reflecting the changes in 28 different metabolites and 8 mineral elements in fully expanded leaves and roots of wheat plants under waterlogging and control treatments. After 12 days of setting up waterlogging, fully expanded leaves and roots were collected from the control and waterlogging treatments. The variance captured by the first and second principal components (PCA1 and PCA2, respectively) is written in brackets. Each arrow represents the loading plots of the dependent variables determined in each organ. The direction and length of the arrows determine the influence of the variables on the different plant organs and treatments.

## Discussion

To date, flooding-related studies in plants have primarily described changes in roots, which are the immediate organs exposed to excess water (Hwang *et al*., 2011). However, it has remained unclear whether and to what extent the integration of metabolomic and transcriptomic modifications through the root-xylem-shoot system are biologically linked and part of the coordinated response of cereal plants to waterlogging. Therefore, our study was focused on investigating the metabolic processes mediated in the root-shoot communication as a systemic response in wheat. In contrast to all previous studies that only examined one time point, the dynamics experiment allowed tracking alanine as the major root-to-shoot translocated and systemically acting metabolite crucial for balancing C/N between roots and shoots.

### Waterlogging affects the physiological status of the plants by alteration of mineral uptake and hormone regulation

Subjecting wheat plants to waterlogging for a period of 12 days resulted in imbalances of nutrient uptake and assimilation. While leaf N concentrations dropped to very low levels of approx. 1%, and root-to-shoot translocation rates became interrupted (Fig. 3A, E), demonstrating a severe deficit in N uptake and allocation. On the other hand, C accumulated in roots as a result of compromised respiration (Fig. 3A, Fig. 4C) indicating substantial disturbance of C and N metabolism under waterlogging. The lower leaf P concentration under waterlogging was associated with lower root-to-shoot translocation rates of P, probably causing a significant upregulation of P transporters in roots to counteract the P deficit (Fig. 3, Supplementary Figure S3B). In some cases, higher P uptake under waterlogging has been related to increased availability of Fe and Mn (Shahandeh *et al*., 1994; Lambers *et al*., 2015; Maranguit *et al*., 2017) as it was also observed in our study. The higher K concentration might be associated to the maintenance of cytosolic K+ homeostasis and channel functionality (Mugnai *et al*., 2011). However, other studies have reported a substantial decline under waterlogging (Smethurst *et al*., 2005; Board 2008). Hormone regulation under waterlogging, including GAs, auxins and cytokinin was also affected as their translocation rate in xylem exudates was highly reduced (Fig. 2). Altogether, these changes were reflected by lower root biomass and root length of stressed plants at early time points, suggesting a rapid decline of energy metabolism in those organs directly exposed to the stress. Initiation of adventitious roots (AR) at later time points compensated for the much shorter seminal roots developed in stressed plants. This phenomenon was observed in wheat plants (Malik *et al*., 2002; Herzog *et al*., 2016) and other species (e.g. maize, ash, willow, Forsythia and *Rumex palustris*) (Bailey-Serres *et al*., 2012). In contrast, aboveground parameters were reduced at the latest time point. Chlorophyll concentration decreased strongly, indicating a reduction in photosynthetic capacity, which has been also observed in other plants species (Bansal and Srivastava 2015). However, other parameters such as the root to shoot ratio remained unaffected. As this latter is an important index for assessing plant health (Agathokleous *et al*., 2019), local response to waterlogging in roots triggered adaptation mechanisms as a systemic response to readjust the root to shoot ratio and thus ensure plant acclimation, which was also reflected in the low penalties of grain development, evaluated in our study (Supplementary Fig. S2).

### Waterlogging triggers systemic response in wheat plants mediated by metabolic and transcriptomic readjustments

Gene expression and metabolite profiling revealed that fermentation processes occurred only in roots where up-regulation of *lactate dehydrogenase* (*LDH*) and *alcohol dehydrogenase* (*ADH*) was observed (Fig. 5). The anaerobic process and the inhibited glycolysis result in higher accumulation of soluble sugars in the roots as demonstrated in our study (Fig. 6). According to Hsu et al. (2011), Arabidopsis roots under waterlogging, reprogram their transcription globally in both roots and shoots. Transcriptional regulation of carbohydrate production has been documented to occur in shoots of waterlogged plants as a systemic response, to which signals must be initiated from waterlogged roots. Reduced rates of anaerobic carbohydrate metabolism occur in waterlogged roots to maintain a low rate of energy provision for extended periods (Gibbs and Greenway 2003). Several genes associated with starch and sucrose metabolism, pentose phosphate pathways, fructose and mannose metabolism, carbon fixation and amino acid metabolism were differentially over-represented in leaves (Fig. 5), suggesting a carbohydrate reprogramming system under waterlogging. Over-expression of *AP2 (APETALA2)/ERF* (Ethylene Responsive Factor)-related genes in both organs suggests that ethylene response transcription factors are prompted to mediate the roots to shoot communication in wheat plants (data not shown). This type of TFs has been repeatedly shown to affect hypoxia responses by modulating ethylene, jasmonic acid (JA) and abscisic acid (ABA) signaling under stress (Lorenzo *et al*., 2003; Zhang *et al*., 2004; Pré *et al*., 2008). The accumulation of sucrose in roots and decrease in leaves was also observed in waterlogged poplar (Kreuzwieser *et al*., 2009). Possible mechanisms to increase sucrose transport to sink organs i.e. roots under ATP limitations, has been described (Gajdanowicz *et al*., 2011; Dreyer *et al*., 2017). The disturbed metabolic pathway in leaves (Fig. 6) might be explained by the differentially expressed genes related to gluconeogenesis and amino acid degradation pathways (Fig. 5) as alternative source of carbons skeletons for glucose biosynthesis and most likely also as nitrogen source (Eastmond *et al*., 2015). Subsequently, amino acid degradation for energy needs might contribute not only to the production of substrates for mitochondrial respiration (Heinemann and Hildebrandt 2021) but also of substrates that undergo gluconeogenesis to form glucose as energy source. Taken altogether, in our study we identified key responsive genes and metabolites that responded in the organ directly exposed to the stress and those ones driven by systemic response as only roots were exposed to waterlogging and leaves were able to adequately respire.

### Alanine is the major root-to-shoot translocated and systemically acting metabolite in waterlogging

Analysis of key metabolites under waterlogging showed that Ala accounted for the highest abundance in roots and xylem exudates (Supplementary Table S1; Fig. 6). However, it decreased in leaves of the same plants. Ala accumulation in waterlogged organs has been associated to pyruvate interconversion to prevent the carbon losses during ethanol fermentation and its diffusion from the cells (Ricoult *et al*., 2006; Miyashita and Good 2008; Schulze *et al*., 2019). This process is catalyzed by *alanine amino transferase* (*ALAT*), which is responsible for the conversion of alanine and oxoglutarate to pyruvate and glutamate in many plant species (Watson *et al*., 1992; Puiatti and Sodek 1999; Rocha, *et al*., 2010; Diab and Limami 2016; Bashar *et al*., 2020). Strong up-regulation of genes encoding *ALAT2* (p<0.001) was found in waterlogged roots in our study, whereas no genes associated with this enzyme were annotated in leaves (Fig. 5, Supplementary Fig. S4). The significance of this enzyme has been discussed in the context of alanine production to regulate pyruvate levels, a product that drives glycolysis, since NAD^+^ is not regenerated during alanine production (De Sousa and Sodek 2003; Rocha *et al*., 2010). Other studies on *Lotus japonicus* and in mutants unable to fix nitrogen via symbiotic interaction with rhizobia showed that Ala accumulation was independent of the nitrogen status of the plant (Rocha *et al*., 2010). These results indicate that alanine metabolism might be a response to prevent pyruvate accumulation and facilitate the continued operation of glycolysis during waterlogging. Subsequently to *ALAT2*, we found strong up-regulation of genes encoding the *alanine glyoxylate aminotransferase 2* (*AGXT2*) (p<0.001) in leaves of stressed plants (Fig. 5, Supplementary Fig. S4). This enzyme is known to participate in the conversion of Ala to pyruvate and Gly using glyoxylate as amino acceptor. To our knowledge, its function in plants has not been well elucidated and it has been restricted to animals, as one of the two structurally distinct types of *AGTs* (Liepman and Olsen 2003). At this point, two questions arise: 1) What is the further use of Alanine in a system of negative nitrogen balance and energy limitation? 2) Is the conversion of Ala to pyruvate in the leaves mediated by AGXT2? The potential benefit of the alanine-pyruvate conversion has been limited to alanine production. In humans, the function of alanine as a “shuttle” to undergo transamination reactions for pyruvate synthesis and the subsequent provision of energy to muscles in form of glucose as a result of gluconeogenesis process is well documented (Petersen *et al*., 2019). However, in plants, little is known about the further contribution of alanine as a carbon-nitrogen source in aboveground organs to replenish *de novo* glucose synthesis and/or nitrogen containing compounds. Based on the observations in the current study in wheat plants under waterlogging, we propose a scheme that integrates a potential mechanism of alanine production in waterlogged roots, its translocation to leaves and further conversion to recycle pyruvate as a carbon source for *de novo* glucose biosynthesis. As illustrated in Fig. 8, limitation of O_2_ concentration in roots caused by waterlogging, results in a partial conversion of pyruvate to alanine via *ALAT2* to prevent excessive carbon loss from the fermentation pathways. Alanine accumulates strongly in the roots as an adaptation response and is translocated to the shoots via xylem exudate. In shoots, alanine and glyoxylate are converted to pyruvate and glycine via *AGXT2*. In the leaves, two possible alternatives may occur: pyruvate is recycled and incorporated into the TCA cycle to contribute to higher net ATP production or pyruvate is incorporated into the gluconeogenesis pathway for *de novo* glucose biosynthesis. The latter can be used immediately in glycolysis to generate additional energy in the form of ATP.

**Fig. 8.**
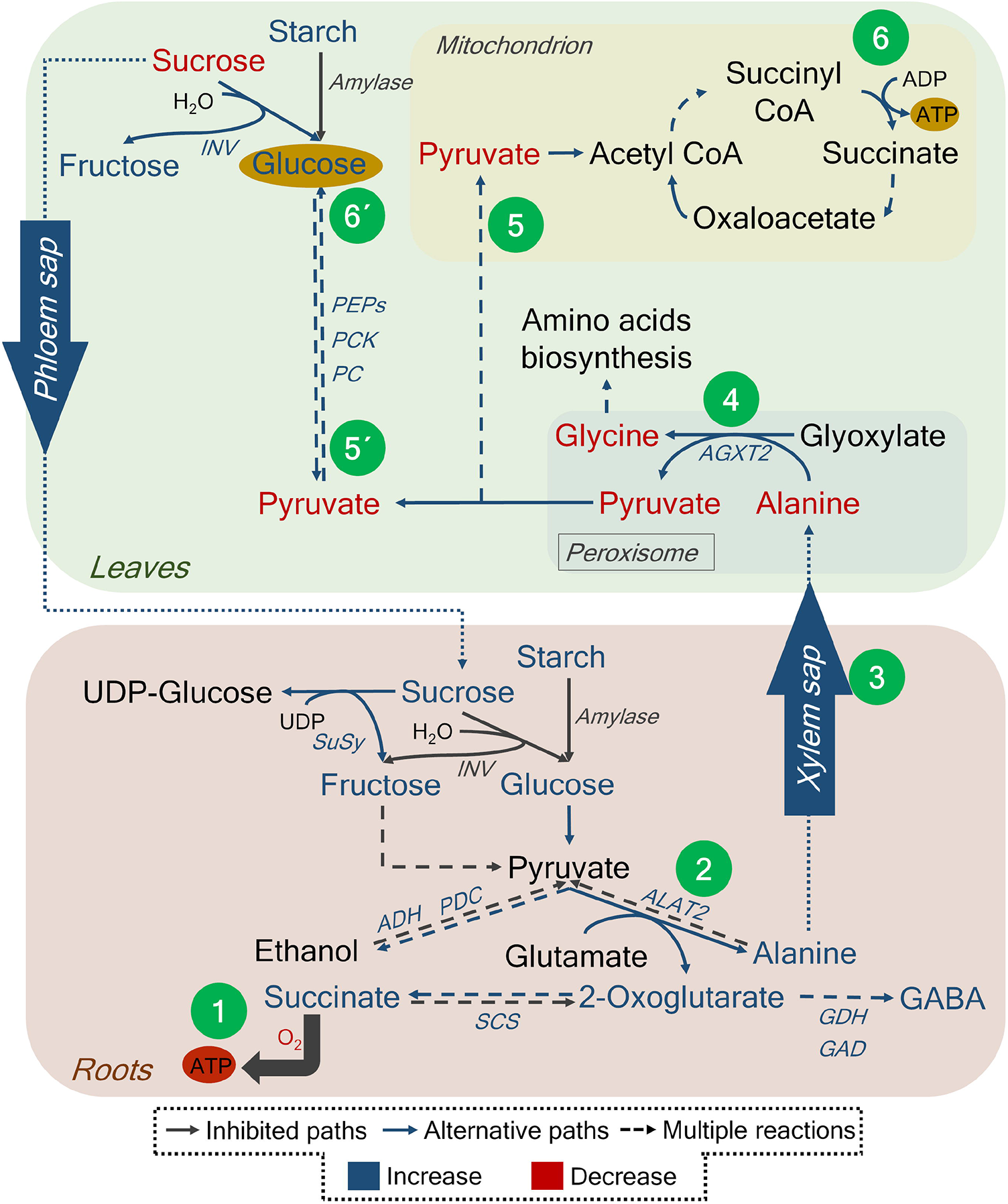
Metabolic adaptations of leaves and roots of wheat plants to cope with waterlogging stress. Under waterlogging, O_2_ concentration decreases and fermentation pathways are activated (1). To prevent the carbon losses due to ethanol production, pyruvate is converted to alanine using alanine amino transferase 2 (ALAT2) (2). The excess alanine is translocated to the shoots via the xylem (3). In shoots, alanine and glyoxylate are converted to pyruvate and glycine using alanine glyoxylate transferase 2 (*AGXT2*) (4). The produced pyruvate can either be recycled and directed to TCA cycle (5) or it is used to synthesize *de novo* glucose via gluconeogenesis pathway (5’, 6’). More glucose can be produced as an energy saving strategy under waterlogging and be further used for additional energy production once the stress is released. *INV*: invertases; *PEPs*: phosphoenolpyruvate synthase; *PCK:* phosphoenolpyruvate carboxykinase; *PC*: pyruvate carboxylase; *AGXT2:* alanine glyoxylate aminotransferase 2; *SuSy*: sucrose synthase; *ADH*: alcohol dehydrogenase; *PDC*: pyruvate decarboxylase; *ALAT2*: alanine aminotransferase 2; *SCS:* succinyl CoA ligase; *GDH:* glutamate dehydrogenase; *GAD:* glutamic acid decarboxylase.

## Summary

Here, we studied the effect of waterlogging in wheat plants at tillering stage based on transcriptome and metabolome approaches. We explored the systemic response of wheat as a whole plant system and the supportive strategy of above-ground organs to meet the energy demand from waterlogged organs. As a consequence of the stress, we showed that levels of plant nutrient uptake and translocation are altered during the exposition to the stress and how this influenced plant biomass and grain-yield parameters. Local stress retarded the metabolism in roots and as consequence, most of the metabolites, including free sugars, organic acids and amino acids were accumulated. Contrary effect was observed in leaves, where there was a higher sugar production but also higher usage and degradation of assimilates to support the unfavorable status of root metabolism. The important role of transport for nutrients, amino acids and hormones in xylem exudates was also explored in this study. Further, we demonstrated how waterlogging affected negatively the translocation rate of important metabolites. However, alanine accumulation in roots and higher translocation in xylem was considered as an important biomarker for the root to shoot communication in wheat plants under waterlogging. This study provided a hypothesis where alanine translocation plays an important role, not only as vehicle for the transport of amino nitrogen but also as a source of carbon skeleton for recycling pyruvate in aboveground organs and thus to contribute to *de novo* glucose biosynthesis as an important metabolic strategy of wheat plants under waterlogging. We also demonstrated that wheat plants are capable to recover after prolonged stress and meet the demand of assimilates to ensure plant acclimation and reproduction. The relevance of this study relies on the basis to characterize the important role of alanine as the major root-to-shoot translocated and systemically acting metabolite crucial for balancing C/N between roots and shoots.

## Acknowledgements

We wish to thank Prof. Andreas Börner for the propagation of the plants and providing the seeds of the spring wheat cultivar used in this study. We thank Nicole Schäfer, Melanie Ruff, Barbara Kettig, Dagmar Böhmert and Heike Nierig (Leibniz Institute of Plant Genetics and Crop Plant Research, IPK) for their excellent assistance during harvesting the plant material, the sample preparation and the biochemical analysis. We also thank Paul Reim (Leibniz Centre for Agricultural Landscape Research, ZALF), Richard Pauwels and Dr. Michael Bitterlich (Leibniz Institute of Vegetable and Ornamental Crops, IGZ) for their great support during experimental setting up and harvesting.

## Credit authorship contribution statement

List of authors contribution: G.C., S.K. and M.R.H. conceived the study and designed the experiments. D.F and N.v.W contributed with the “dynamics experiment” design. G.C. performed all experiments. G.C. and M.R.H performed the biochemical assays and analyzed the data, D.F. and Y.T.M helped with the data processing. N.v.W contributed with the scientific methodology and data interpretation. Y.T.M performed the elemental analysis and hormone measurement. G.C. wrote the manuscript. D.F., S.K., N.v.W. and M.R.H. revised critically the manuscript.

## Data availability

The RNA seq.dataset is available on the server of the Institute of Plant Genetics and Crop Plant Research (IPK).

## Appendix Supporting information

**Table S1**. Amino acids concentration in leaves, roots and translocation rate in xylem sap of wheat plants exposed to 12 days waterlogging at tillering stage

The difference between the mean of paired groups was analyzed using student’s T-test.

*p<0.05; **p<0.01; ***p<0.001; ns: not significant; na: not analyzed; nq: not quantified).

**Table S2**. Top soil physicochemical properties of the soil used in this study

The numbers represent the mean of three replicates and the standard deviation.

**Table S3**. Characterization of the soil used in this study after 12 days of exposition to waterlogging

The difference between the mean of paired groups was analyzed using student’s T-test. (p≤ 0.05; n=8) (**p<0.01; ***p<0.001;).

**Fig. S1**. Experimental design to evaluate the effect of waterlogging on wheat plants at tillering stage.

**Fig. S2:** Effect of waterlogging during tillering stage on grain yield-related parameters at ripening stage. (A) Total spike weight, (B) spike number, (C) total grain weight per plant and (D) number of grains per plants. After 12 days of setting up waterlogging, the excess of water was removed allowing the plants to recovered their metabolism and growth performance. At ripening stage, spikes and grains were harvested from plants of control, recovered from waterlogging and not-recovered from waterlogging treatments. The line in the middle of the boxes are plotted as the median and box limits indicate the 25^th^ and 75^th^ percentiles. Whiskers extend to 1.5 times the interquartile range from the 25^th^ and 75^th^ percentiles. Normal distribution of the data was checked using Shapiro-Wilk’s normality test. To determine the difference between paired groups, one-way ANOVA was performed. Different letters indicate significant differences between the treatments based on Tukey’s HSD. In all cases, 95% confidence interval (p≤ 0.05, n=8) was considered.

**Fig. S3**. Relative expression pattern of genes expressed in fully expanded leaves and roots. (A) Relative expression of genes related to nitrogen uptake and translocation, (B) phosphate transporters, (C) potassium transporters and channels and (D) iron transporters. In all cases, genes were grouped according to their transcript abundance at day 12 after setting up the waterlogging treatment. Genes were selected considering a threshold >2 in absolute value and padjust < 0.05.

**Fig. S4**. Relative expression pattern of genes encoding transaminases expressed in fully expanded leaves and roots. Relative expression of genes related to (A) ALAT2- and (B): AGXT2. In all cases, genes were grouped according to their transcript abundance at day 12 after setting up the waterlogging treatment. Genes were selected considering a threshold >2 in absolute value and padjust < 0.05. The line in the middle of the boxes are plotted as the median and box limits indicate the 25^th^ and 75^th^ percentiles. Whiskers extend to 1.5 times the interquartile range from the 25^th^ and 75^th^ percentiles.

**Fig. S5**. Water retention curve to estimate the water holding capacity of the soil used in this study.

## References

Agathokleous E, Belz R. G, Kitao M, Koike T, Calabrese EJ. 2019. Does the root to shoot ratio show a hormetic response to stress? An ecological and environmental perspective. Journal of Forestry Research 30(5), 1569–1580.

Aguirre A. 2020. Quantification of the impact of enhanced urea uptake in cereal grain crops on growth, nitrogen metabolism, yield formation and grain quality. PhD Thesis, Martin-Luther-Universität Halle-Wittenberg. http://dx.doi.org/10.25673/37297. Accessed March 2021.

Arbona V, Gómez-Cadenas A. 2008. Hormonal modulation of citrus responses to flooding. Journal of Plant Growth Regulation 27(3), 241–250.

Bailey-Serres J, Fukao T, Gibbs DJ, Holdsworth MJ, Lee SC, Licausi F, Perata P, Voesenek LACJ, van Dongen JT. 2012. Making sense of low oxygen sensing. Trends in Plant Science 17(3), 129–138.

Bailey-Serres J, Lee SC, Brinton E. 2012. Waterproofing crops: effective flooding survival strategies. Plant Physiology 160(4), 1698–1709.

Bansal R, Srivastava J. 2015. Effect of waterlogging on photosynthetic and biochemical parameters in pigeonpea. Russian Journal of Plant Physiology 62(3), 322–327.

Bashar KK, Tareq MZ, Islam M. S. 2020. Unlocking the mystery of plants’ survival capability under waterlogging stress. Plant Science Today 7(2), 142–153.

Beier S, Marella NC, Yvin JC, Hosseini SA, von Wirén N. 2022. Silicon mitigates potassium deficiency by enhanced remobilization and modulated potassium transporter regulation. Environmental and Experimental Botany 198, 104849.

Board J. 2008. Waterlogging effects on plant nutrient concentrations in soybean. Journal of Plant Nutrition 31(5), 828–838.

Bradford KJ, Hsiao TC, Yang SF. 1982. Inhibition of ethylene synthesis in tomato plants subjected to anaerobic root stress. Plant Physiology 70(5), 1503–1507.

Cui J, Davanture M, Zivy M, Lamade E, Tcherkez G. 2019. Metabolic responses to potassium availability and waterlogging reshape respiration and carbon use efficiency in oil palm. New Phytologist 223(1), 310–322.

De Sousa C, Sodek L. 2003. Alanine metabolism and alanine aminotransferase activity in soybean (Glycine max) during hypoxia of the root system and subsequent return to normoxia. Environmental and Experimental Botany 50(1), 1–8.

Diab H, Limami AM. 2016. Reconfiguration of N metabolism upon hypoxia stress and recovery: roles of alanine aminotransferase (AlaAT) and glutamate dehydrogenase (GDH). Plants 5(2), 25.

Dreyer I, Gomez-Porras JL, Riedelsberger J. 2017. The potassium battery: a mobile energy source for transport processes in plant vascular tissues. New Phytologist 216(4), 1049–1053.

Eastmond PJ, Astley HM, Parsley K, Aubry S, Williams BP, Menard GN, Craddock CP, Nunes-Nesi A, Fernie AR, Hibberd JM. 2015. Arabidopsis uses two gluconeogenic gateways for organic acids to fuel seedling establishment. Nature Communications 6(1), 1–8.

Eggert K, von Wirén N. 2017. Response of the plant hormone network to boron deficiency. New Phytologist 216(3), 868–881.

Ellis MH, Dennis ES, James Peacock W. 1999. Arabidopsis roots and shoots have different mechanisms for hypoxic stress tolerance. Plant Physiology 119(1), 57–64.

Francioli D, Cid G, Kanukollu S, Ulrich A, Hajirezaei MR, Kolb S. 2021. Flooding Causes Dramatic Compositional Shifts and Depletion of Putative Beneficial Bacteria on the Spring Wheat Microbiota. Frontiers in Microbiology 12.

Gajdanowicz P, Michard E, Sandmann M, Rocha M, Corrêa LGG, Ramírez-Aguilar SJ, Gomez-Porras JL, González W, Thibaud JB, van Dongen JT. 2011. Potassium (K+) gradients serve as a mobile energy source in plant vascular tissues. Proceedings of the National Academy of Sciences 108(2), 864–869.

Ghaffari MR, Shahinnia F, Usadel B, Junker B, Schreiber F, Sreenivasulu N, Hajirezaei MR. 2016. The metabolic signature of biomass formation in barley. Plant and Cell Physiology 57(9), 1943–1960.

Gibbs J, Greenway H. 2003. Mechanisms of anoxia tolerance in plants. I. Growth, survival and anaerobic catabolism. Functional Plant Biology 30(1), 1–47.

Heinemann B, Hildebrandt TM. 2021. The role of amino acid metabolism in signaling and metabolic adaptation to stress-induced energy deficiency in plants. Journal of Experimental Botany 72(13), 4634–4645.

Herzog M, Striker GG, Colmer TD, Pedersen O. 2016. Mechanisms of waterlogging tolerance in wheat–a review of root and shoot physiology. Plant, Cell & Environment 39(5), 1068–1086.

Hsu FC, Chou MY, Peng HP, Chou SJ, Shih MC. 2011. Insights into hypoxic systemic responses based on analyses of transcriptional regulation in Arabidopsis. PLoS One 6(12), e28888.

Hwang JH, Lee MO, Choy YH, Ha-Lee YM, Hong CB, Lee DH. 2011. Expression profile analysis of hypoxia responses in Arabidopsis roots and shoots. Journal of Plant Biology 54(6), 373–383.

Irfan M, Hayat S, Hayat Q, Afroz S, Ahmad A. 2010. Physiological and biochemical changes in plants under waterlogging. Protoplasma 241(1), 3–17.

León J, Cruz-Castillo M, Gayubas B. 2020. The hypoxia-reoxygenation stress in plants. Journal of Experimental Botany 72, 16.

Kreuzwieser J, Hauberg J, Howell KA, Carroll A, Rennenberg H, Millar AH, Whelan J. 2009. Differential response of gray poplar leaves and roots underpins stress adaptation during hypoxia. Plant Physiology 149(1), 461–473.

Lambers H, Hayes PE, Laliberte E, Oliveira RS, Turner BL. 2015. Leaf manganese accumulation and phosphorus-acquisition efficiency. Trends in Plant Science 20(2), 83–90

Liepman AH, Olsen LJ. 2003. Alanine aminotransferase homologs catalyze the glutamate: glyoxylate aminotransferase reaction in peroxisomes of Arabidopsis. Plant Physiology 131(1), 215–227.

Lorenzo O, Piqueras R, Sánchez-Serrano JJ, Solano R. 2003. ETHYLENE RESPONSE FACTOR1 integrates signals from ethylene and jasmonate pathways in plant defense. The Plant Cell 15(1), 165–178.

Lothier J, Diab H, Cukier C, Limami AM, Tcherkez G. 2020. Metabolic responses to waterlogging differ between roots and shoots and reflect phloem transport alteration in Medicago truncatula. Plants 9(10), 1373.

Malik AI, Colmer, TD, Lambers H, Setter TL, Schortemeyer M. 2002. Short-term waterlogging has long-term effects on the growth and physiology of wheat. New Phytologist 153(2), 225–236.

Maranguit D, Guillaume T, Kuzyakov Y. 2017. Effects of flooding on phosphorus and iron mobilization in highly weathered soils under different land-use types: Short-term effects and mechanisms. Catena 158, 161–170.

Miyashita Y, Good AG. 2008. Contribution of the GABA shunt to hypoxia-induced alanine accumulation in roots of Arabidopsis thaliana. Plant and Cell Physiology 49(1), 92–102.

Mugnai S, Marras AM, Mancuso S. 2011. Effect of hypoxic acclimation on anoxia tolerance in Vitis roots: response of metabolic activity and K+ fluxes. Plant and Cell Physiology 52(6), 1107–1116.

Mustroph A, Barding-Jr GA, Kaiser KA, Larive CK, Baley-Serres J. 2014. Characterization of distinct root and shoot responses to low-oxygen stress in A rabidopsis with a focus on primary C-and N-metabolism. Plant, Cell & Environment 37(10), 2366–2380.

Oliveira HC, Freschi L, Sodek L. 2013. Nitrogen metabolism and translocation in soybean plants subjected to root oxygen deficiency. Plant Physiology and Biochemistry 66, 141–149.

Orzechowski S, Socha-Hanc J, Paszkowski A. 1999. Alanine aminotransferase and glycine aminotransferase from maize (Zea mays L.) leaves. Acta Biochimica Polonica 46(2), 447–457.

Petersen KF, Dufour S, Cline GW, Shulman GI. 2019. Regulation of hepatic mitochondrial oxidation by glucose-alanine cycling during starvation in humans. The Journal of Clinical Investigation 129(11), 4671–4675.

Pré M, Atallah M, Champion A, De Vos M, Pieterse CM, Memelink J. 2008. The AP2/ERF domain transcription factor ORA59 integrates jasmonic acid and ethylene signals in plant defense. Plant Physiology 147(3), 1347–1357.

Puiatti M, Sodek L. 1999. Waterlogging affects nitrogen transport in the xylem of soybean. Plant Physiology and Biochemistry 37(10), 767–773.

Reggiani R. 1999. Amino acid metabolism under oxygen deficiency. Phytochemistry 2, 171–174.

Reggiani R, Nebuloni M, Mattana M, Brambilla I. 2000. Anaerobic accumulation of amino acids in rice roots: role of the glutamine synthetase/glutamate synthase cycle. Amino acids 18(3), 207–217.

Ricoult C, Echeverria LO, Cliquet JB, Limami AM. 2006. Characterization of alanine aminotransferase (AlaAT) multigene family and hypoxic response in young seedlings of the model legume Medicago truncatula. Journal of Experimental Botany 57(12), 3079–3089.

Roca M, Chen K, Pérez-Gálvez A. 2016. 6. Chlorophylls. In: Carle R, Schweiggert RM eds. Handbook on natural pigments in food and beverages. Industrial Applications for Improving Food Color (pp. 125-158).

Rocha M, Licausi F, Araújo WL, Nunes-Nesi A, Sodek L, Fernie AR, Van Dongen JT. 2010. Glycolysis and the tricarboxylic acid cycle are linked by alanine aminotransferase during hypoxia induced by waterlogging of Lotus japonicus. Plant Physiology 152(3), 1501–1513.

Rocha M, Sodek L, Licausi F, Hameed MW, Dornelas MC, Van Dongen JT. 2010. Analysis of alanine aminotransferase in various organs of soybean (Glycine max) and in dependence of different nitrogen fertilisers during hypoxic stress. Amino acids 39(4), 1043–1053.

Schulze ED, Beck E, Buchmann N, Clemens S, Müller-Hohenstein K, Scherer-Lorenzen M. 2019. Oxygen deficiency In: Schulze ED, Beck E, Buchmann N, Clemens S, Müller-Hohenstein k, Scherer-Lorenzen M. Plant Ecology, 143–164.

Shahandeh H, Hossner L, Turner F. 1994. Phosphorus relationships in flooded rice soils with low extractable phosphorus. Soil Science Society of America Journal 58(4), 1184–1189.

Smethurst CF, Garnett T, Shabala S. 2005. Nutritional and chlorophyll fluorescence responses of lucerne (Medicago sativa) to waterlogging and subsequent recovery. Plant and Soil 270(1), 31–45.

Tognetti VB, Zurbriggen MD, Morandi EN, Fillat MF, Valle EM, Hajirezaei MR, Carrillo N. 2007. Enhanced plant tolerance to iron starvation by functional substitution of chloroplast ferredoxin with a bacterial flavodoxin. Proceedings of the National Academy of Sciences 104(27), 11495–11500.

Tula S, Shahinnia F, Melzer M, Rutten T, Gómez R, Lodeyro AF, von Wirén N, Carrillo N, Hajirezaei MR. 2020. Providing an additional electron sink by the introduction of cyanobacterial flavodiirons enhances growth of A. thaliana under various light intensities. Frontiers in plant science 11, 902.

Watson NR, Peschke VM, Russell DA, Sachs MM. 1992. Analysis of alanine: 2-oxoglutarate aminotransferase isozymes in maize. Biochemical genetics 30(7), 371–383.

Zhang H, Huang Z, Xie B, Chen Q, Tian X, Zhang X, Zhang H, Lu X, Huang D, Huang R. 2004. The ethylene-, jasmonate, abscisic acid-and NaCl-responsive tomato transcription factor JERF1 modulates expression of GCC box-containing genes and salt tolerance in tobacco. Planta 220(2), 262–270.

